# Atypical NMDA Receptors Limit Synaptic Plasticity in the Adult Ventral Hippocampus

**DOI:** 10.1101/2022.10.05.510966

**Authors:** Emily P. Hurley, Bandhan Mukherjee, Lisa Fang, Jocelyn R. Barnes, Firoozeh Nafar, Michiru Hirasawa, Matthew P. Parsons

## Abstract

N-methyl-D-aspartate receptors (NMDARs) assemble as functionally diverse heterotetramers. Incorporation of the GluN3A subunit into NMDARs alters conventional NMDAR properties by reducing both magnesium sensitivity and calcium permeability. GluN1 together with GluN3A can also form functional receptors that lack a glutamate binding site and instead serve as excitatory glycine receptors (eGlyRs). GluN3A expression is high in early development but naturally declines to low levels in most brain regions by adulthood. Interestingly, GluN3A expression remains elevated in the CA1 of the adult ventral hippocampus (VH), but not in the dorsal hippocampus (DH). The DH and VH are now well-understood to play very different functional roles, with the DH being primarily involved in cognitive functions and the VH in emotional processing. Why GluN3A persists in the adult VH, and the impact its presence has on glutamatergic neurotransmission in the VH is currently unknown. Here, we show that GluN3A remains elevated both at synaptic and extrasynaptic locations in the adult VH, assembling as GluN1/GluN2/GluN3A NMDARs with reduced magnesium sensitivity, as well as GluN1/GluN3A eGlyRs. By comparing various synaptic properties in the DH and VH of wild-type (WT) and GluN3A knockout (KO) mice, we demonstrate that GluN3A persistence in the VH attenuates glutamate release, limits postsynaptic calcium influx through NMDARs, and reduces the magnitude of NMDAR-dependent long-term potentiation. In comparison, GluN3A KO had relatively little effect on these same properties in the DH. In all, our data demonstrate that GluN3A persistence in the VH represents a key modulator of VH excitability and therefore may play a central role in emotional processing.

## Introduction

N-methyl-D-aspartate (NMDA) receptors (NMDARs) are ionotropic glutamate receptors that have essential roles in synaptic development, transmission, and plasticity (Paoletti et al., 2013). The NMDAR subunit GluN3A, encoded by the *Grin3A* gene, may be the most perplexing of the receptor’s subunits. NMDARs assemble as functional heterotetramers with two obligatory GluN1 subunits, and activation of classical GluN1/GluN2 NMDARs can induce canonical forms of synaptic plasticity known to play essential roles in learning and memory (Luscher & Malenka, 2012; Maren & Baudry, 1995). In contrast, the incorporation of GluN3A subunits yields unconventional NMDARs with atypical biophysical properties that seemingly counteract the functions of classical GluN1/GluN2 NMDARs. For example, GluN3A-containing NMDARs have a reduced Mg^2+^ sensitivity (Roberts et al., 2009; Sasaki et al., 2002), and restriction of Ca^2+^ permeability (Pérez-Otaño et al., 2016). GluN3A can also function as a dominant-negative regulator of NMDAR-mediated synaptic plasticity (Pérez-Otaño et al., 2016) and limit NMDAR-dependent activation of plasticity genes such as *Bdnf* and *Arc* (Chen et al., 2020). The GluN3A subunit can be incorporated into NMDARs to assemble as GluN1/GluN2/GluN3A NMDARs or GluN1/GluN3A excitatory glycine receptors (eGlyRs). The latter has been identified in the adult medial habenula (Otsu et al., 2019), and juvenile CA1 pyramidal neurons (Grand et al., 2018). More recently, eGlyRs have been identified in somatostatin-positive interneurons in the adult cortex, and pyramidal neurons in the adult basolateral amygdala, where they are mainly located extrasynaptically to sense ambient glycine and generate tonic excitatory currents (Bossi et al., 2022). Although rarer than their postsynaptic counterparts, GluN3A-containing NMDARs have also been observed at presynaptic terminals where they can regulate transmitter release (Bouvier et al., 2015; Larsen et al., 2011, 2014; Savtchouk et al., 2019).

GluN3A expression predominates during a narrow window of postnatal development in many brain regions, including the CA1 hippocampus (Henson et al., 2010; Pérez-Otaño et al., 2016; Wong et al., 2002). Throughout most of the central nervous system, GluN3A expression declines in the second and third postnatal week and remains low into adulthood (Pérez-Otaño et al., 2016). However, it has become clear that GluN3A does not decline equally in all brain regions. In the adult brain, GluN3A expression is virtually absent in the CA1 subregion of the dorsal hippocampus (DH), yet high GluN3A levels persist into adulthood in CA1 of the ventral hippocampus (VH) (Cembrowski et al., 2016; Murillo et al., 2021). It has become increasingly-recognized that the hippocampus differs considerably along its longitudinal axis, with the DH being primarily responsible for spatial cognitive functions such as learning and memory, while the VH is more heavily involved in emotional and stress processing (Fanselow & Dong, 2010; Strange et al., 2014). The functional consequences of GluN3A persistence in the adult VH are unknown.

It is well-established the DH and VH exhibit differences in short and long-term plasticity (Colgin et al., 2004; Maggio & Segal, 2007a); the DH undergoes robust long-term potentiation (LTP), whereas the VH exhibits a significantly weaker LTP (Maggio & Segal, 2007b). Prolonging GluN3A expression beyond the physiological window at CA3-CA1 was shown to reduce the magnitude of LTP (Roberts et al., 2009), consistent with the idea of GluN3A being a dominant-negative regulator of classical NMDAR function (Pérez-Otaño et al., 2016). In this study, we aimed to determine the purpose of the natural GluN3A retention in the adult VH. Therefore, the goals of the current study are to first, determine GluN3A location, whether it exist at synaptic sites, extrasynaptic sites, or both? Second, how does GluN3A incorporate into NMDARs in the adult VH? Does GluN3A exist as GluN1/GluN2/GluN3A NMDARs or as GluN1/GluN3A eGlyRs? Lastly, does GluN3A presence in the adult VH impact synaptic structure or function?

## Methods

### Animals

Two to four-month old male and female GluN3A KO (Jax strain 029974) and wildtype (WT) B6129SF2/J (Jax strain 101045) mice were used in the present experiments. Mice had *ad libitum* access to food and water and were housed on a 12 h:12 h light:dark cycle. All procedures followed the guidelines of the Canadian Council on Animal Care and were approved by Memorial University’s Institutional Animal Care Committee.

### Subcellular fractionation

Whole brains were dissected from GluN3A KO or WT mice and placed in chilled phosphate buffer saline for 2 minutes. For each hemisphere, the hippocampus was rolled out and DH and VH samples collected and flash frozen in liquid nitrogen. Subcellular fractionation and western blotting protocols were performed as previously described (Milnerwood et al., 2010). The GluN3A antibody used was obtained from Millipore Sigma (cat #07-356).

### Stereotaxic surgery

For the duration of the surgical procedure, mice were anesthetized with isoflurane. Viral injections of 1 μl hSyn.iGluSnFr.WPRE.SV40 (Addgene, plasmid #98929-AAV1) for glutamate imaging or 1 μl pENN.AAV.CamKII.GCaMP6f.WPRE (Addgene, plasmid #100834-AAV1) for calcium imaging were administered directly into the hippocampus. Adeno-associated virus was injected using a Neuros 7002 Hamilton syringe coupled to an infusion pump (Pump 11 Elite Nanomite; Harvard Apparatus), which allowed a constant infusion rate of 2 nl/s. The syringe was left in place for an additional 5 min. The following coordinates from bregma were used for DH infusions: 2.6 mm posterior, 2.3 mm lateral, and 1.1 to 1.3 mm ventral to the brain surface. The following coordinates from bregma were used for VH infusions: 3.3 mm posterior, 3.0 mm lateral, and 3.3 mm ventral to the brain surface.

### Acute slice preparation

Two to four weeks after adeno-associated virus injection, mice were anesthetized by isoflurane inhalation, and brains were quickly removed and placed in oxygenated ice-cold slicing solution containing the following (in mm): 125 NaCl, 2.5 KCl, 25 NaHCO_3_, 1.25 NaH_2_PO_4_, 2.5 MgCl_2_, 0.5 CaCl_2_, and 10 glucose. Hippocampal slices (350 μm) were obtained using a Precisionary Compresstome VF-310-0Z. Slices were transferred to a holding chamber containing oxygenated ACSF, the same as slicing solution except containing 1 mm MgCl_2_ and 2 mm CaCl_2_, for recovery (>60 min) before experimentation. For whole-cell patching acute slices were sliced in NMDG ACSF slicing solution as described (Ting et al., 2018), recovered in HEPES ACSF recovery solution with sodium spike in every 5 minutes for one hour before transferred to standard ACSF for 45 minutes (Ting et al., 2018).

### Electrophysiology

Electrophysiological recordings of LTP were performed as previously described (Barnes et al., 2020). Acute hippocampal slices were placed in a recording chamber and were left for a minimum of 10 min before electrode placement. Slices were visualized with an Olympus BX51 microscope. Field potentials were recorded with a glass electrode in CA1 stratum radiatum and stimulated with a glass electrode placed in the Schaffer collaterals. The resistance of the stimulating and recording pipettes were between 1-2 MΩ when filled with ACSF. LTP was induced with standard high frequency stimulation (HFS) protocol of 1 X HFS, 100 Hz for 1 second. Recordings continued for 30 minutes after LTP induction. LTP magnitude was quantified by averaging the fEPSP slope throughout the last 5 min of recording. Percent potentiation was expressed as the percentage increase in the average fEPSP slope compared with baseline. All data were collected and analyzed using pClamp10 software (Molecular Devices). For Mg^2+^ sensitivity experiments field responses were evoked using two pulses with 10ms interval. Responses were recorded in 0.1mM MgCl_2_ ACSF and 20μM DNQX solution before switching to 1mM MgCl_2_ ACSF and 20μM DNQX solution. The percent decrease of the fEPSP response was measured to determine inhibition by Mg^2+^.

### Whole-cell patching

To perform whole-cell patch clamp experiments, slices were transferred to the recording chamber and perfused with ACSF at a rate of 1-2mL/min and bubbled with 95% O_2_/5% CO_2_. The chamber was maintained at 27-30°C. Pyramidal cells within the CA1 layer were visualized using infrared differential interference contrast optics (DM LFSA, Leica Microsystems, ON, Canada). Glass recording pipettes had a tip resistance of 3-5 MΩ when filled with internal solution (composition in mM: 123 K gluconate, 2 MgCl_2_, 1 KCl, 0.2 EGTA, 10 HEPES, 5 Na_2_ATP, 0.3 Na_2_GTP, and 2.7 biocytin). Glass recording pipettes were used for puffing experiments and had a resistance of 3-5 MΩ when filled with glycine (10mM). Puffing pipettes were positioned downstream of ACSF flow, approximately 20 μm away from the soma, ventral to the cell being patched. Cells were held at −70mV and a 500ms puff was applied using a Picospritzer III (Parker Hannifin, NH, USA) in the presence of picrotoxin (GABA_A_ receptor antagonist; 50 μM), strychnine (glycine receptor antagonist; 20 μM), D-APV (NMDA receptor antagonist; 50 μM), DNQX (AMPA and kainite receptor antagonist; 10 μM), CGP-78608 (GluN1/GluN3A potentiator; 1μM) (Grand et al., 2018). Data were recorded using Multiclamp 700B amplifier and pClamp 10 software (Molecular Devices).

### Golgi staining

Golgi staining was performed using the sliceGolgi kit and protocol (Bioenno Lifesciences, cat# 003760). Whole brains were dissected and placed in Golgi fixative overnight. Once fixed, 100 μm dorsal or ventral hippocampal slices were obtained using Precisionary Compresstome. Slices were stained following the protocol and stains from the sliceGolgi Kit and mounted on gelatin-coated slides and cover slipped using Permount Mounting media. Z-stack images of 5-10 stacks were taken using a 63X objective and Zeiss Axio microscope. For image analysis, a 10 μm section of a secondary dendritic branch of a CA1 pyramidal apical dendrite was selected. Length and width measurements were determined for each dendritic spine using ImageJ software. Spines were classified based on length and width ratios, as described previously (Risher et al., 2014).

### Real-time imaging of glutamate and calcium biosensors

Following recovery, slices were transferred to a recording chamber with oxygenated ACSF was continuously perfused at a flow rate of 1.5–2 ml/min. For glutamate imaging, the Schaffer collateral pathway was stimulated by controlling an Iso-flex stimulator as previously described (Barnes et al.,2020). iGluSnFR responses were evoked by stimulating the Schaffer collateral pathway with a single burst of HFS and iGluSnFR fluorescence was excited by a LED (Prior Scientific, Lumen 300) passed through a 460–490 nm bandpass filter. Clampex software was used to synchronize the LED, stimulus isolator, and image acquisition. Images were acquired using Andor Solis software. For calcium imaging, GCaMP6f baseline fluorescence was measured in Mg^2+^ free ACSF with 0.5 μM TTX followed by a 2-minute bath application of Mg^2+^ free ACSF with 0.5 μM TTX, 10 μM NMDA and 20 μM glycine before switching back to Mg^2+^ free ACSF with 0.5 μM TTX alone. Clampex software was used to synchronize the LED, stimulus isolator, and image acquisition. Images were acquired using Andor Solis software every 10 seconds for the duration of the experiment.

### Statistics

All data are represented as mean ± SEM. All statistical analyses were conducted using GraphPad Prism. The statistical tested used include unpaired t-test and two-way ANOVA. The specific statistical test used is listed within the results. p values of at least 0.05 were considered significant. Reported n values indicate the number of acute slices used, and for every dataset, a minimum of 3 mice were used per group.

## Results

### GluN3A expression is elevated at synaptic and extrasynaptic sites in the adult VH and assembles as GluN1/GluN2/GluN3A NMDARs and GluN1/GluN3A eGlyRs

Expression of GluN3A has been previously shown to be high in the adult VH (Cembrowski et al., 2016; Murillo et al., 2021). Electron microscopy localization studies from the juvenile brain suggest that most GluN3A exists extrasynaptically (Pérez-Otaño et al., 2016). First, we sought to determine the subcellular localization of GluN3A subunits in the adult VH. Through subcellular fractionation we isolated synaptic and extrasynapic GluN3A protein expression in 2-4-month-old wild-type (WT) mice. Indeed, compared to the adult DH, GluN3A expression is elevated in the adult VH. Interestingly, we show that VH GluN3A is elevated at both extrasynaptic and synaptic sites (Figure 1A-C, n = 4 animals, synaptic WT DH vs. VH p = 0.0063, extrasynaptic WT DH vs. VH p = 0.0031, unpaired t-test). When we used this same protocol on tissue from 2-4-month-old GluN3A KO (KO) mice, the GluN3A protein was not detected either synaptically or extrasynaptically (Figure 1A-B), verifying both the antibody and the GluN3A KO mouse. Thus, GluN3A is present at both synaptic and extrasynaptic sites in the adult VH.

**Figure 1:**
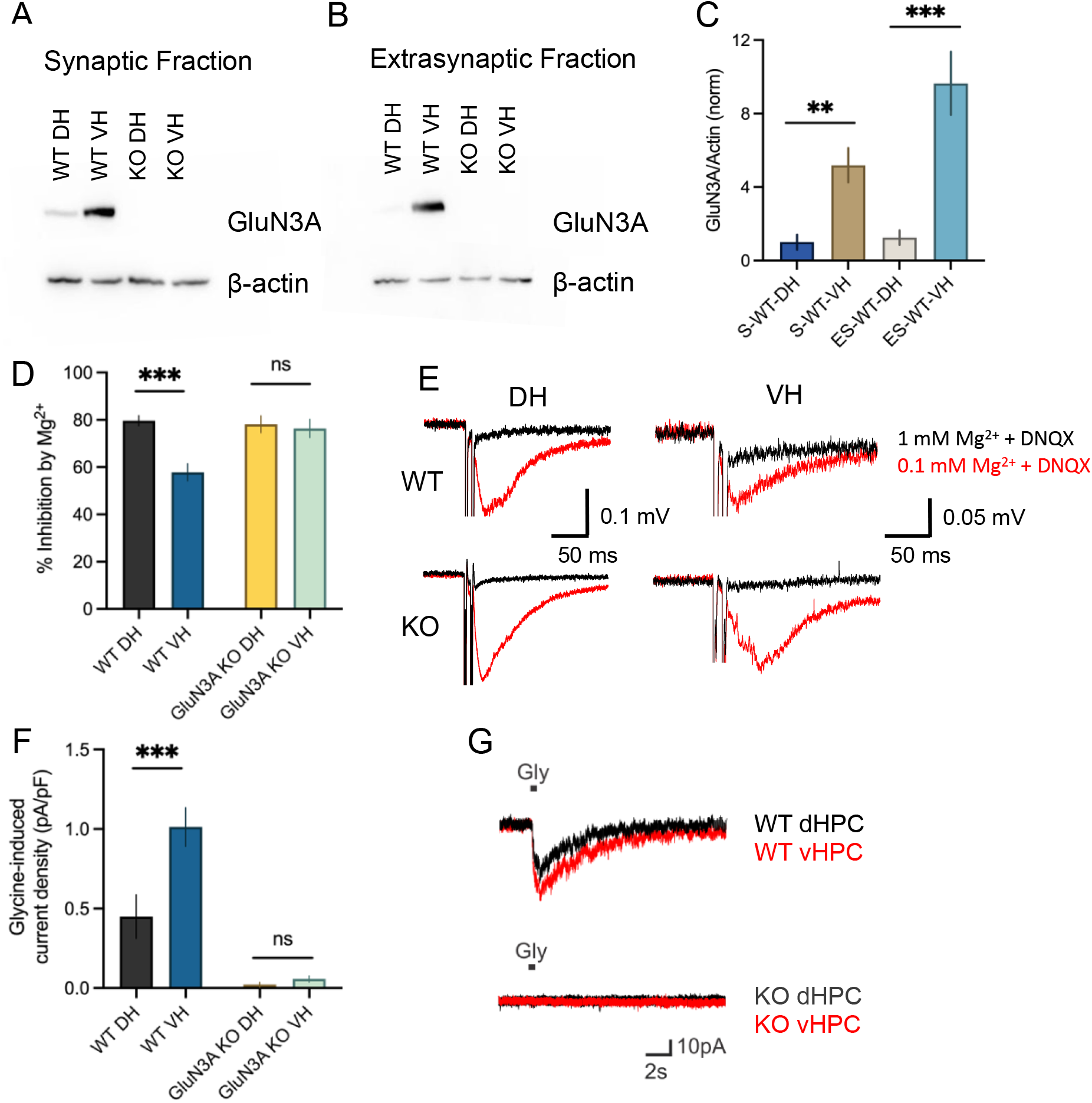
The adult wild-type VH has higher expression of GluN3A-containing NMDARs that assemble in GluN1/GluN2/GluN3A and eGlyR configurations. **A-B)** Western blot of synaptic and extrasynaptic GluN3A expression in DH (dorsal hippocampus) and VH (ventral hippocampus) of adult wild-type (WT) and GluN3A KO (KO) mice. **C)** Bar graph of synaptic (S) and extrasynaptic (ES) GluN3A expression in WT DH and VH (normalized). **D)**% inhibition by 1mM Mg^2+^ in CA1 Schaffer collaterals. **E)** Representative traces in 0.1 mM Mg^2+^ ACSF and 20 μM DNQX (red trace) and 1mM Mg^2+^ ACSF and 20 μM DNQX (black trace). **F)** Current density of glycine-induced responses (pA/pF). **G)** Representative traces of glycine-induced current. Black bar indicates time of glycine puff. Error bars represent s.e.m., ** p<0.01, *** p<0.001.

Incorporation of the GluN3A subunit into NMDARs can result in the formation of GluN1/GluN2/GluN3A triheteromeric NMDARs that bind glutamate (Hansen et al., 2014; Pérez-Otaño et al., 2016) or GluN1/GluN3A eGlyRs that do not bind glutamate (Bossi et al., 2022; Grand et al., 2018; Pérez-Otaño et al., 2016). To determine how GluN3A-containing NMDARs assemble in the WT VH we first tested Mg^2+^ sensitivity, as GluN1/GluN2/GluN3A NMDARs have reduced sensitivity to Mg^2+^ blockade (Roberts et al., 2009; Sasaki et al., 2002). To test NMDAR sensitivity to Mg^2+^ in either the DH or VH, NMDAR fEPSPs were recorded in low Mg^2+^ ACSF in the presence of 20 μM DNQX to block AMPA and kainite receptors. Glutamate release was evoked by electrical stimulation of the Schaffer collaterals. Following a stable NMDAR-mediated fEPSP response to Schaffer collateral stimulation, the bath solution was switched to high Mg^2+^ ACSF and we quantified the percent inhibition of the NMDAR fEPSP by Mg^2+^. In WT mice, the DH exhibited high sensitivity to Mg^2+^ as expected, with approximately 80% of the NMDAR fEPSP being blocked by 1 mM Mg^2+^. In contrast, the VH exhibited significantly reduced Mg^2+^ sensitivity in WT mice (Figure 1D-E, n = 10 slices over 5-6 animals, p <0.001, Tukey’s test, 2-way ANOVA). The observed reduction in Mg^2+^ sensitivity in the VH was dependent on GluN3A presence as both the DH and VH were highly sensitive to Mg^2+^ in GluN3A KO mice (Figure 1D-E n = 8 slices over 4 animals, p = 0.988, Tukey’s test, 2-way ANOVA). These data suggest that at least some of the GluN3A present in the adult VH assembles as triheteromeric GluN1/GluN2/GluN3A NMDARs that are responsive to synaptically-evoked glutamate release and exhibit reduced sensitivity to Mg^2+^ blockade.

As GluN3A subunits can also assemble as GluN1/GluN3A receptors that do not bind glutamate and are an eGlyR, we next quantified glycine currents in the DH and VH using whole-cell patch clamp electrophysiology. CA1 pyramidal cells were recorded in acute slices from either the DH or VH, and glycine (10 μM) was applied via puff pipette in the presence of multiple receptor antagonists (see methods) and 1 μM CGP-78609 to reduce eGlyR desensitization. The WT mice, excitatory glycine current density was significantly higher in the VH compared to the DH, suggesting a higher presence of eGlyRs in the VH (Figure 1F-G, n = 11 cells for DH, 15 cells for VH, over 4 animals, p = 0.01, Tukey’s test, 2-way ANOVA). Glycine responses were not observed whatsoever in either the DH or VH of GluN3A KO mice (Figure 1F-G, n = 10 cells for DH, 10 cells for VH over 4 animals per genotype, p = 0.996, Tukey’s test, 2-way ANOVA), confirming that the responses we observed in WT mice were GluN3A-dependent eGlyR currents. Together, these results demonstrate that GluN3A expression is high in the adult VH, located at both synaptic and extrasynaptic locations, and appears to assemble as both GluN1/GluN2/GluN3A NMDARs with reduced Mg^2+^ sensitivity and as GluN1/GluN3A eGlyRs.

### GluN3A KO enhances LTP in the adult VH

Research on the mechanism underlying postsynaptic NMDAR-mediated LTP is most often conducted at the CA3-CA1 synapse in the hippocampus; thus, much of what we know regarding NMDAR-dependent LTP is based on observations at this specific connection in the brain. However, the magnitude of CA3-CA1 LTP can vary tremendously along the hippocampal longitudinal axis; LTP is weak in the VH compared to intermediate and dorsal regions of the hippocampus (Maggio & Segal, 2007b). In addition, short-term plasticity also differs at the CA3-CA1 synapses in the DH and VH, with paired-pulse ratios (PPRs) being significantly reduced in the VH compared to the DH (Maggio & Segal, 2007a). The reasons underlying this differential capacity for both short- and long-term plasticity is poorly understood. As GluN3A overexpression has been demonstrated to reduce the magnitude of LTP (Roberts et al., 2009), and GluN3A is elevated in the adult VH, we hypothesized that the naturally occurring retention of GluN3A in the adult VH may act as a molecular brake on synaptic plasticity in this region. As a measure of short-term plasticity, paired-pulse ratios (PPR) were measured at 50, 100, and 250 ms interpulse intervals. Compared to the WT DH, the WT VH showed a lower PPR at these interpulse-intervals (Figure 2A, n =20 slices over 7-8 animals. pulse-interval p <0.001, regional p <0.001, 2-way ANOVA). In addition, and again consistent with previous reports, LTP induced by 1X HFS (100Hz, 1 sec) in the WT VH was significantly weaker than in the DH (Figure 2B and F, n = 25 slices for DH, 16 slices for VH over 9-10 animals, p <0.001, Tukey’s test, 2-way ANOVA). Short- and long-term plasticity was also quantified in the DH and VH of age-matched GluN3A KO mice. Like WT mice, the VH showed a lower PPR at 50, 100 and 250 ms intervals than DH in GluN3A KO mice (Figure 2C, n = 18 slices for DH, 20 slices for VH over 7-8 animals, pulse-interval p <0.001, regional p <0.001, 2-way ANOVA), suggesting that the blunted short-term facilitation in the VH is independent of GluN3A subunits. It is worth noting in the DH, GluN3A KO significantly increased PPR at the 50 ms interval, suggesting some role of GluN3A in presynaptic short-term plasticity (Figure 2E, p = 0.049, Tukey’s test, 2-way ANOVA). When we examined long-term plasticity, we found that GluN3A KO had a profound effect on LTP in the VH. In mice lacking GluN3A, DH LTP was not significantly affected, but VH LTP was now found to exceed that observed in the DH (Figure 2D and F, n = 12 slices for DH, 11 slices for VH over 4-5 animals, p = 0.023, Turkey’s test, 2-way ANOVA), and VH LTP was considerably higher in GluN3A KOs compared to VH LTP in WT mice (Figure 2F, p<0.001, Tukey’s test, 2-way ANOVA). Therefore, the relatively high expression of GluN3A that persists into adulthood in the VH does indeed act as a brake on synaptic plasticity.

**Figure 2:**
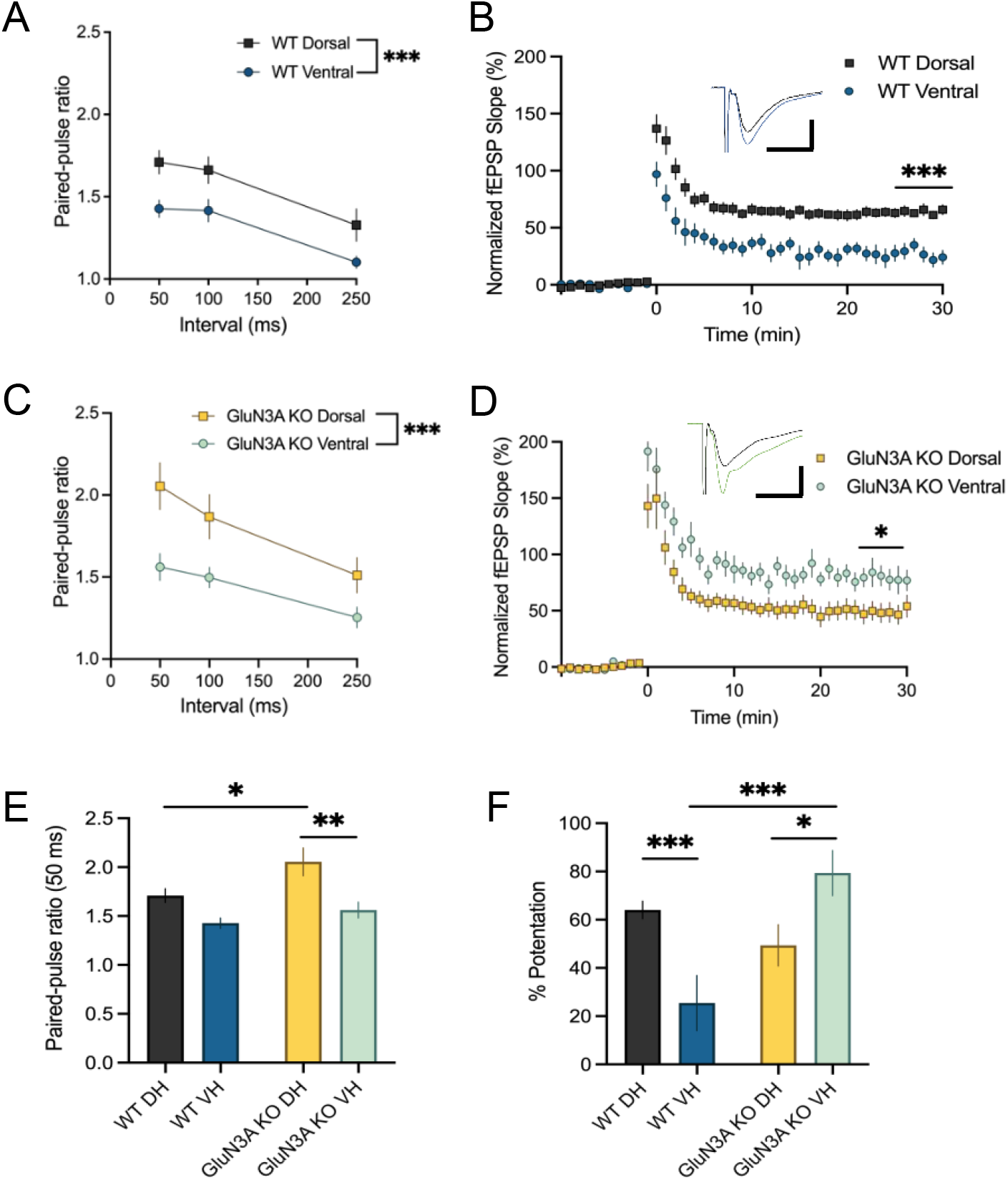
GluN3A knockout enhances LTP in the adult VH. **A)** Paired-pulse ratios for interpulse-intervals of 50, 100 and 250 ms in 2–4-month-old WT DH (black) and WT VH (blue). **B)** Graph of normalized fEPSP (excitatory postsynaptic potential) slope (%) following high frequency stimulation (HFS) (HFS; applied at time 0) in 2-4-month-old WT DH and WT VH. Representative field potential traces recorded from VH CA1 stratum radiatum in response to single pulse before and 30 min after HFS (black; baseline recording, blue; potentiated response). Scale bars: 200 μV, 20 ms. **C)** Paired-pulse ratios for interpulse-intervals of 50, 100 and 250 ms in 2–4-month-old GluN3A KO DH (yellow) and GluN3A KO VH (green). Representative field potential traces recorded from CA1 stratum radiatum in response to single pulse Schaffer collateral stimulation. **D)** Graph of normalized fEPSP (%) following HFS (HFS; applied at time 0) in 2-4-month-old GluN3A KO DH and VH. Representative field potential traces recorded from VH CA1 stratum radiatum in response to single pulse before and 30 min after HFS (black; baseline recording, green; potentiated response). Scale: 400 μV, 20 ms. **E)** Bar graph representing the paired-pulse ratios at 50 ms. **F)** Bar graph representing % potentiation measured 25-30 min after HFS. Error bars represent s.e.m. * p<0.05, ** p<0.01, *** p<0.001.

### GluN3A persistence in the VH does not impact spine maturity

NMDARs are principal mediators of synaptic plasticity, an activity-dependent process that can alter structural plasticity via postsynaptic dendritic spines. As GluN3A overexpression has been shown to reduce spine maturation, spine density and stability (Kehoe et al., 2014; Roberts et al., 2009), it is possible that GluN3A retention in the adult VH maintains dendritic spines in an immature state. Thus, we imaged Golgi-stained CA1 pyramidal dendrites to determine if GluN3A expression impacts basal spine density and/or morphology. The maturation of dendritic spines serves as a structural measure of synaptic strength, where stable mushroom spines have strong synaptic connections, and filipodia and thin spines have a higher rate of turnover and exhibit weaker synaptic connections (Risher et al., 2014). To investigate structural synaptic strength, we categorized dendritic spines by type as a measure of maturity. In WT mice, we found no significant difference between overall spine density in the DH and VH (Figure 3B, n = 24 images over 3 animals per region, p = 0.274, Tukey’s test, 2-way ANOVA). Similarly, spine density was also similar between the DH and VH in GluN3A KO mice (Figure 3B, n = 16 images over 2 animals per region, p = 0.620, Tukey’s test, 2-way ANOVA). In WT and GluN3A KO mice, there was no significant difference in mushroom spine density in the DH or VH (Figure 3C, n = 24 images over 3 animals per region, regional p = 0.253, genotype p = 0.058, interaction p = 0.290, 2-way ANOVA). Similarly, in both WT and GluN3A KO mice, the DH and VH had a similar density of thin spines (Figure 3D, n = 24 images over 3 animals per region, regional p = 0.634, genotype p = 0.038, interaction p = 0.899, 2-way ANOVA), although we observed a significant reduction in thin spines in GluN3A KO mice compared to WT, regardless of the region examined (2-way ANOVA genotype p = 0.038). Regarding long-thin spines, we found more long-thin spines in the VH compared to the DH, although this was independent of GluN3A expression (Figure 3E, n = 24 images over 3 animals per region, regional p = 0.05, genotype p = 0.109, interaction p = 0.760, 2-way ANOVA). Lastly, there were no significant differences in stubby spine density between the WT and GluN3A KO mice in either the DH or VH (Figure 3F, n = 24 images over 3 animals per region, regional p = 0.113, genotype p = 0.368, interaction p = 0.484, 2-way ANOVA). Overall, these data demonstrate that GluN3A persistence in the VH does not negatively impact spine maturation.

**Figure 3:**
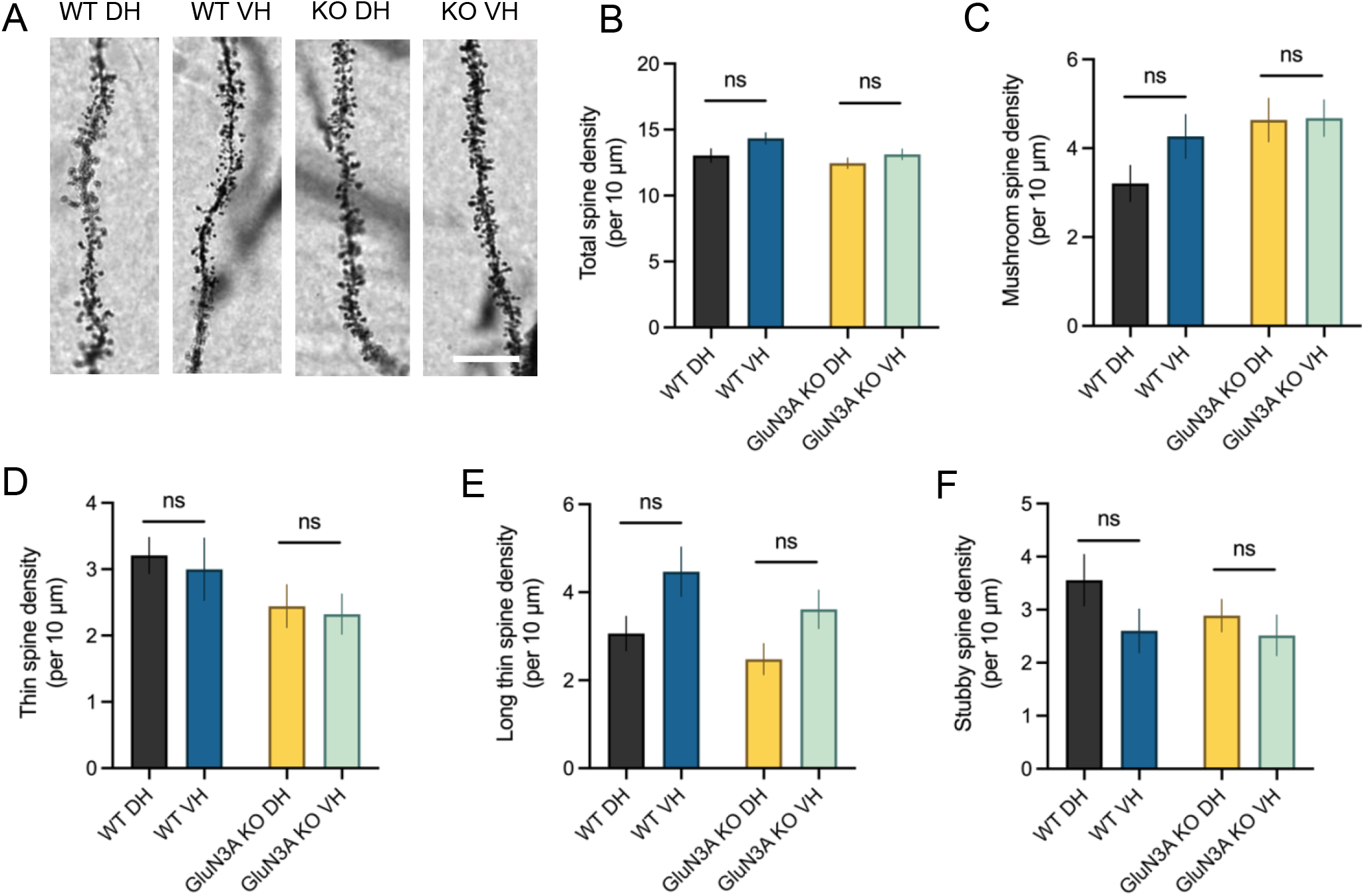
Dendritic spines of the VH are just as mature as the DH. **A)** Representative images of CA1 pyramidal neurons in 2-4-month-old wild-type and GluN3A KO mice (Scale= 10 μm). **B)** Total spine density per 10 μm segment. **C)** Mushroom spine density per 10 μm segment. **D)** Thin spine density per 10 μm segment. **E)** Long-thin spine density per 10 μm segment. **F)** Stubby spine density per 10 μm segment. Error bars represent s.e.m. ns, not significant.

### GluN3A knockout boosts glutamate release to a greater extent in the VH

Although rarer than their postsynaptic counterparts, GluN3A has been observed at many presynaptic terminals where they can regulate and enhance evoked transmitter release (Bouvier et al., 2015; Larsen et al., 2011, 2014; Savtchouk et al., 2019). As GluN3A is expressed at high levels in the adult VH, we asked whether putative presynaptically-located GluN3A could potentially alter the glutamate release during LTP induction during HFS. To quantify the release amount of glutamate released during HFS, WT and GluN3A KO mice were injected into the DH or VH with the glutamate biosensor iGluSnFR (Marvin et al., 2013) and acute hippocampal slices were obtained two to four weeks later. iGluSnFR fluorescence was measured in the stratum radiatum while HFS was applied to the Schaffer collaterals. We quantified the area under the curve of HFS-evoked iGluSnFR transients, thereby capturing the entire iGluSnFR profile to serve as a measure of total glutamate accumulation during LTP induction. Interestingly, while not significant, we see trending differences in extracellular glutamate in the WT VH compared to the DH in response to HFS, as measured by the area under the curve, where the VH tends to accumulate more glutamate than the DH (Figure 4C, n= 8 slices over 4-5 animals, p = 0.0791, Tukey’s test, 2-way ANOVA). In GluN3A KO mice, iGluSnFR responses were increased overall (Figure 4C, genotype p = 0.0004, 2-way ANOVA), with a particularly large response now observed in the VH (Figure 4C, n=8 slices over 4-5 animals, p <0.0001, Tukey’s test, 2-way ANOVA). GluN3A KO effectively doubled the size of iGluSnFR responses to HFS in the VH when compared to the WT (Figure 4C, n=8 slices over 4-5 animals, p = 0.0007, Tukey’s test, 2-way ANOVA). Thus, GluN3A presence reduced glutamate release during HFS, with a greater suppressant effect in the VH compared to the DH. Thus, it is possible that GluN3A KO boosts VH LTP in part by enhancing glutamate release during LTP induction.

**Figure 4:**
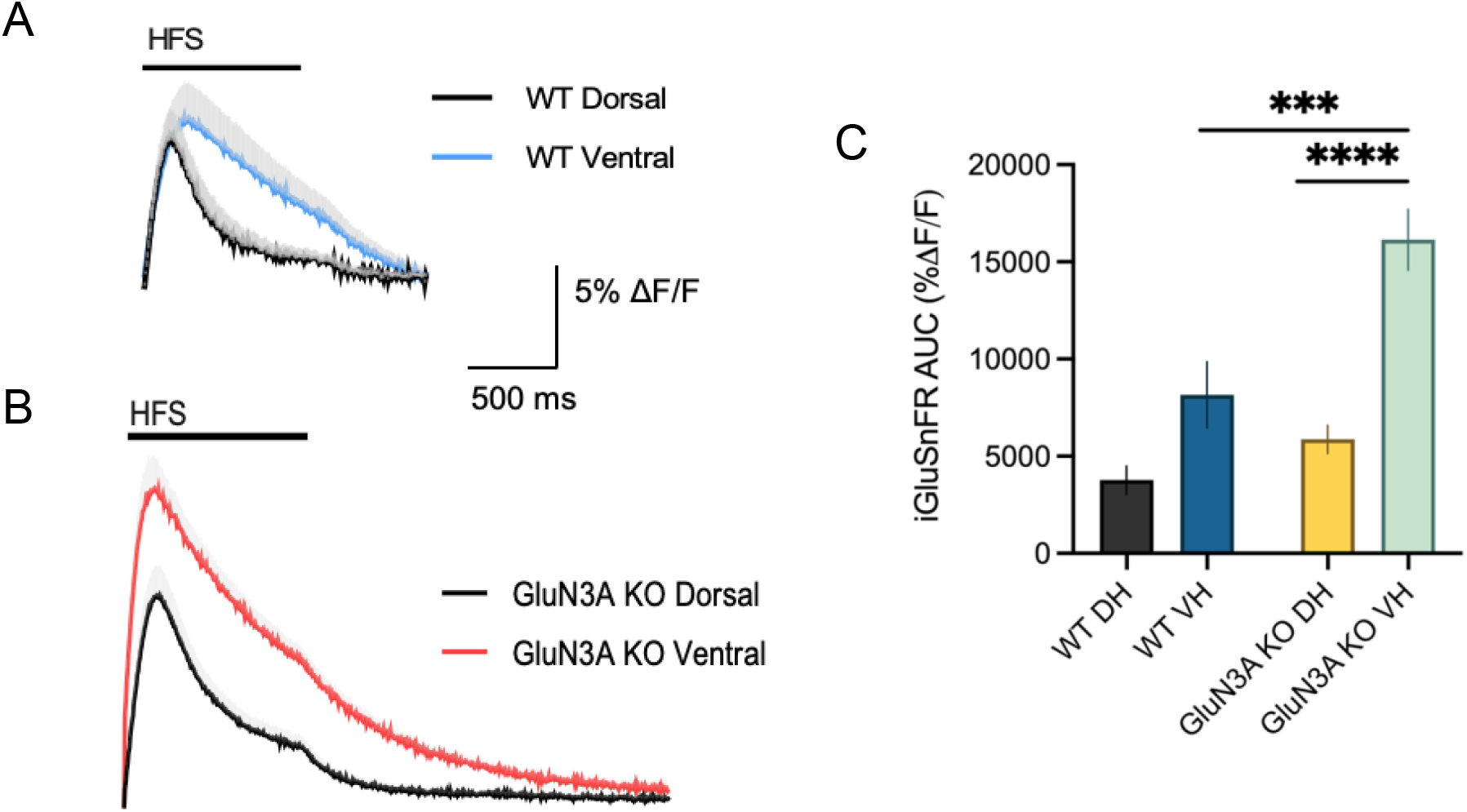
GluN3A knockout enhances glutamate release in the VH. **A)** Representative trace of glutamate response recorded from CA1 stratum radiatum during HFS. Shown are the WT DH (black) and the WT VH (blue) responses. **B)** Representative trace of glutamate response recorded from CA1 stratum radiatum during HFS. Shown are the GluN3A KO DH (black) and the GluN3A KO VH (red) responses. **C)** iGluSnFR area under the curve (AUC). Error bars represent s.e.m. *** p<0.001, **** p<0.0001.

### GluN3A knockout boosts postsynaptic Ca^2+^ responses to exogenous NMDA to a greater extent in the adult VH

Most literature reports that GluN3A incorporation decreases Ca^2+^ permeability through NMDARs (Chatterton et al., 2002; Madry et al., 2010; Sasaki et al., 2002). As Ca^2+^ influx directly plays a role in synaptic plasticity by triggering postsynaptic signalling pathways (Kawamoto et al., 2012), we asked whether GluN3A persistence in the VH suppresses Ca^2+^ responses to NMDAR activation. WT and GluN3A KO mice were injected with the calcium biosensor GCaMP6f, and acute hippocampal slices were obtained two-four weeks later. GCaMP6f fluorescence was measured before, during and after a 2-minute bath application of ACSF with 0.5 μM TTX, 10 μM NMDA and 20 μM glycine. GCaMP6f responses were confirmed to be NMDAR dependent as no increase in GCaMP6f fluorescence was observed when NMDARs were blocked with dAPV (Figure 5C). There were no significant differences in the GCaMP6f area under the curve between the WT DH and VH (Figure 5A and E, n= 14 slices over 4-5 animals, p = 0.995, Tukey’s test, 2-way ANOVA), nor in peak GCaMP6f response between WT DH and VH (Figure 5A and F, n= 14 slices, p =0.998, Turkey’s test, 2-way ANOVA). However, in mice lacking GluN3A, GCaMP6f responses to NMDA in the VH were dramatically elevated compared to WT (Figure 5E, n= 14 slices over 4-5 animals for WT, n= 7 over 4-5 animals for GluN3A KO, p <0.0001, Tukey’s test, 2-way ANOVA). GCaMP6f peak responses were also significantly larger in the VH of GluN3A KO mice compared to WT (Figure 5F, p <0.0001, Tukey’s test, 2-way ANOVA). Note that GluN3A KO also boosted DH Ca^2+^ peak response, but to a lesser extent than that observed in the VH (Figure 5E-F, p = 0.0447, Tukey’s test, 2-way ANOVA). These data demonstrate that GluN3A presence limits Ca^2+^ influx through NMDARs in the hippocampus, with a greater suppressant effect in the VH compared to the DH.

**Figure 5:**
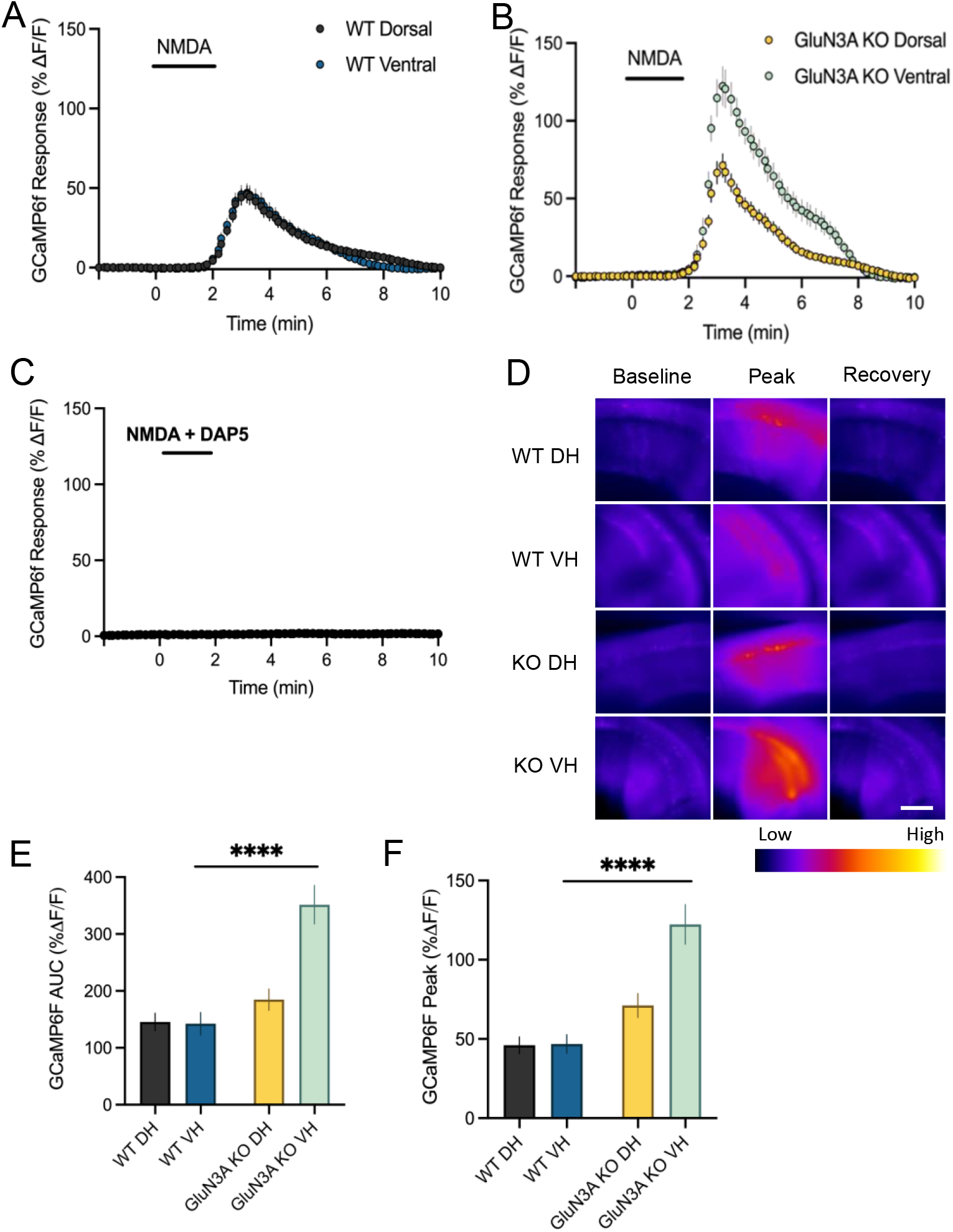
GluN3A knockout enhances calcium responses to exogenous NMDA in the VH. **A)** GCaMP6f response from CA1 stratum radiatum during a 2-minute application of 10 μM NMDA and 20 μM glycine in WT DH and VH. **B)** GCaMP6f response from CA1 stratum radiatum during a 2-minute application of 10 μM NMDA and 20 μM glycine in GluN3A KO DH and VH. **C)** Complete lack of GCaMP6f response in CA1 stratum radiatum during a 2-minute application of 10 μM NMDA, 20 μM glycine and 50 μM DAP5 to block NMDARs in WT DH **D)**Heat map showing response during baseline, peak and recovery (Scale = 1μm). **E)** GCaMP6f area under the curve (AUC). **F)** Peak GCaMP6f response (%ΔF/F). Error bars represent s.e.m. **** p<0.0001.

## Discussion

In the present study, we provide subcellular fractionation and electrophysiological data to demonstrate clear evidence that in the adult VH, GluN3A expression is elevated where it acts to limit the magnitude of synaptic plasticity. We further show that GluN3A-containing NMDARs assemble in both GluN1/GluN2/GluN3A and eGlyR configurations in the hippocampus. Using electrophysiology and fluorescent biosensor imaging, we demonstrate that in the VH, GluN3A knockout boosts glutamate release during trains of electrical stimulation, dramatically enhances postsynaptic responsiveness to exogenous NMDA, and enhances NMDAR-dependent LTP to a level that exceeds that observed in the DH. Thus, GluN3A retention alters various properties in the adult VH that serve to limit synaptic potentiation; how such an arrangement is beneficial for the emotional roles ascribed to the VH remains to be seen and is of interest for future studies.

### Putative locations of GluN1/GluN2/GluN3A NMDARs and eGlyRs

We report that GluN3A is highly expressed at synaptic and extrasynaptic sites in the adult VH (Figure 1A-B), however the putative locations of the GluN1/GluN2/GluN3A NMDARs and the eGlyRs are unclear. A recent paper from Bossi et al. 2022 demonstrated that rather than concentrating at the synapse, these eGlyRs are evenly distributed diffusely throughout the dendrite (Bossi et al., 2022). From our exploration of these two GluN3A receptor configurations in this study, we show a reduced sensitivity to Mg^2+^ to synaptically-evoked currents. Due to the nature of this experiment, synaptic locations would be primarily affected, and as a result, the reduced Mg^2+^ sensitivity detected would likely orginate from synaptic NMDARs. In addition, we use whole-cell patching to evoke large responses to glycine puff application. The experimental design here would likely affect both synaptic and extrasynaptic locations. While further work needs to be done, it is possible that the GluN1/GluN2/GluN3A NMDARs are located mainly at the synapse, where they can respond to synaptically-released glutamate, while the eGlyRs express diffusely throughout the dendrite to respond to ambient glycine. Putative differences in the locations of these two receptor types are supported by known differences in subunit architecture. Classical NMDARs have strong attachment within the PSD, this is due to the PDZ-binding motif that is present only on the C-terminal tail of GluN1 and GluN2 subunits (Bard et al., 2010; Eriksson et al., 2007). Since GluN3A subunits lack this PDZ-binding motif (Eriksson et al., 2007), perhaps GluN3A-containing NMDARs have lesser attachment to PSDs than classical NMDARs, and for this reason have been previously shown to predominate at extrasynaptic locations (Pérez-Otaño et al., 2016).

### Functional characterization of GluN1/GluN2/GluN3A NMDARs and eGlyRs

GluN1/GluN2/GluN3A NMDARs are activated by glutamate and glycine, whereas eGlyRs are activated by glycine alone. Glycine binding to the GluN3A ligand binding domain (LBD) triggers channel opening and activation, whereas glycine binding to GluN1 LBD causes rapid desensitization. To evoke glycine-activated currents, several studies have used the compound CGP-78609, a competitive antagonist with a preference for the GluN1 glycine binding site over GluN3A (Grand et al., 2018; Yao & Mayer, 2006). Application of CGP-78608 prevents glycine binding on GluN1, thus reducing desensitization which unmasks large glycine-induced currents mediated by GluN1/GluN3A eGlyRs. Such glycine-activated currents in the presence of CGP-78608 have been recorded in CA1 pyramidal neurons in young (P8-P12) wild-type mice (Grand et al., 2018) and more recently, in adult wild-type SST-interneurons and BLA pyramidal neurons, but not in GluN3A KO mice (Bossi et al., 2022) To the best of our knowledge, the present paper is the first to show evidence for eGlyRs in principle neurons in the adult hippocampus. Recently, eGlyRs were shown to induce a tonic inward current to low concentrations of ambient glycine (Bossi et al., 2022), and interestingly, CA1 neurons have been previously reported to be more depolarized in the VH compared to the DH (Malik et al., 2016). Thus, tonic excitatory glycine currents mediated by eGlyRs may play a major role in VH excitability.

### Natural expression of GluN3A in the adult VH does not impact dendritic spines

GluN3A overexpression has been shown prevent synapse maturation (Pérez-Otaño et al., 2016). Here, we investigated whether the persistence of GluN3A in the adult VH negatively impacts dendritic spine density and/or morphology. Through examining dendritic spines in the DH and VH of WT and GluN3A KO mice, we found that GluN3A persistence in the VH does not seem to alter basal spine morphology. Our findings may seem to contrast previous studies that have demonstrated that GluN3A overexpression altered the dendrite spine profile (Kehoe et al., 2014; Roberts et al., 2009). However, it is important to note that these studies where done in transgenic mouse models that had prolonged GluN3A expression (Roberts et al., 2009), and in DIV18 cell culture neurons that overexpressed GluN3A (Kehoe et al., 2014). Thus, the naturally occurring GluN3A expression in the VH does not appear to have the same effect of preventing spine maturation. Furthermore, our data suggest that GluN3A’s ability to suppress LTP in the VH is unlikely to be due to any GluN3A-induced alterations in basal spine morphology.

### GluN3A presence reduces glutamate release and postsynaptic Ca^2+^ responses in the adult VH

As glutamate is the predominate excitatory neurotransmitter in the central nervous system, activating ionotropic glutamate receptors to induce synaptic plasticity (Luscher & Malenka, 2012), we investigated the role of GluN3A presence on glutamate release to determine whether this subunit can impact the amount of glutamate released during HFS in the adult hippocampus. Analysis of the area under the curve of iGluSnFR responses to HFS in the DH and VH of WT mice revealed that the VH tends to accumulate more glutamate than the DH in general, and that this regional difference is also observed in GluN3A KO mice. However, the magnitude of the iGluSnFR response, regardless of the region examined, is elevated in the absence of GluN3A, suggesting that GluN3A does function to some extent in limiting glutamate accumulation. Interestingly, the largest boost to glutamate release was observed in the VH following GluN3A KO, consistent with the much higher expression of GluN3A in the VH compared to the DH. It is of interest for future studies to determine whether the observed effects of GluN3A KO on glutamate release is due to presynaptically-located GluN3A-containing NMDARs.

Previous studies have demonstrated that GluN3A incorporation reduces Ca^2+^ influx through NMDARs. GluN1/GluN2/GluN3A NMDARs were reported to have a 10-fold lower Ca^2+^ permeability than classical NMDARs (Sasaki et al., 2002), whereas eGlyRs are Ca^2+^ impermeable (Chatterton et al., 2002; Madry et al., 2010). In the present study, we found that the postsynaptic calcium response to NMDA bath application was similar in the DH and VH, despite the VH having significantly elevated GluN3A expression. However, knocking out GluN3A increased the Ca^2+^ responses to NMDA in both the DH and VH, but to a much greater extent in the VH. These results suggest that in the absence of GluN3A, NMDAR-mediated Ca^2+^ influx is substantially higher in the VH compared to the DH. Whether the elevated Ca^2+^ responsiveness in the VH is due to a higher surface expression of conventional NMDARs, increased Ca^2+^ permeability, and/or increased single channel conductance remains to be seen. Nonetheless, GluN3A retention in the adult VH clearly serves to dramatically supress this otherwise large Ca^2+^ response. In all, our biosensor imaging data demonstrate a role of GluN3A to suppress glutamate release and reduce postsynaptic Ca^2+^ influx in the adult VH.

### GluN3A persistence and LTP in the adult VH

We show that GluN3A KO boosts glutamate release and Ca^2+^ preferentially in the VH. We also show that GluN3A KO boosts LTP in the VH but not in the DH. Thus, it is tempting to speculate that the VH normally exhibits low levels of LTP, relative to the DH, due to a GluN3A-mediated suppression of glutamate release and postsynaptic Ca^2+^ responses. However, it is important to note that in WT mice, neither glutamate release nor postsynaptic Ca^2+^ responses to NMDA were reduced in the VH compared to the DH. It is possible that, for reasons unknown at present, the VH must reach a higher threshold to exhibit robust plasticity, and by removing GluN3A we unmasked a glutamate and Ca^2+^ response sufficient to surpass this threshold. It will be of interest for future studies to determine how GluN3A surface levels are regulated in the VH, as the sensitivity of the VH to activity-dependent plasticity appears to be highly dictated by GluN3A expression. In addition to its influence over glutamate release and Ca^2+^ entry, the C-terminal of the GluN3A subunit interacts with many different proteins with known roles in synaptic plasticity, including protein phosphatase 2A (PP2A). PP2A is a major serine-threonine phosphatase highly expressed in the central nervous system (Strack 1998), that like NMDARs contribute to synaptic transmission and plasticity (Chan & Sucher, 2001). The PP2A protein physically associates with GluN3A via the c-terminal domain and this interaction enhances PP2A activity (Chan & Sucher, 2001; Kehoe et al., 2013; Pérez-Otaño et al., 2016). This GluN3A-PP2A interaction serves to dephosphorylate GluN1 at the serine 897 residue and can impair LTP (Chan & Sucher, 2001; Kehoe et al., 2013; Li et al., 2009). Thus, it is possible that high levels of GluN3A in the adult VH may keep PP2A in its active state to dephosphorylate GluN1 at serine 897 to limit LTP.

### Significance of the GluN3A persistence in the adult VH

We show that GluN3A remains elevated at both synaptic and extrasynaptic locations in the adult VH. We hypothesized that prolonged GluN3A expression acts as a brake on VH plasticity. Indeed, in wild type mice LTP was significantly lower in the VH compared to the DH. In contrast, GluN3A KO mice exhibited strong VH LTP that exceeded the magnitude of LTP in the DH, therefore GluN3A presence negatively impacts LTP. We speculate that the adult VH naturally expresses high GluN3A to help limit synaptic plasticity in this region to serve the different functions of the DH and VH. The DH is responsible for primarily cognitive functions, such as spatial learning, memory, and navigation. Therefore, low GluN3A in the DH allows for robust synaptic plasticity to support spatial learning and memory functions. However, the VH is primarily involved in emotional and stress processing, sharing bidirectional connectivity with the BLA, known to regulate emotions and anxiety (Chan & Sucher, 2001). High levels of GluN3A in the VH may serve to prevent the overexcitation of this brain region that may enhance emotional responses, stress, and anxiety behaviours. In other words, GluN3A persistence may limits VH plasticity in adulthood to serve a protective role to prevent overexcitation of anxiety and fear circuitry. It is of interest for future studies to determine whether GluN3A knockdown specifically in the adult VH enhances behavioural measures of fear and anxiety.

## Acknowledgments

This work was supported by funding from an Early Career Capacity Building Grant, Brain Canada, and the Azrieli Foundation. Thank you to the Animal Care staff at MUN for the assistance with animal housing and monitoring.

